# Variation in the Gut Microbiota of Common Marmosets: Differences with Colony of Origin and Integration

**DOI:** 10.1101/2020.08.31.276733

**Authors:** Rachel E. Cooper, Lisa M. Mangus, Jessica Lynch, Kayla Schonvisky, Justin R. Wright, Christopher J. McLimans, Hoi Tong Wong, Jeremy Chen See, Regina Lamendella, Joseph L. Mankowski

**Affiliations:** Department of Molecular and Comparative Pathobiology, Johns Hopkins University, Baltimore, MD, USA; Department of Biomedical Engineering, Johns Hopkins University, Baltimore, MD, USA; WrightLabs LLC., Huntingdon, PA, USA

## Abstract

Characterization of the gut microbiome may aid understanding and management of natural and experimental disease states in research animals, thereby promoting reproducibility. In this study, the rectal bacterial communities of three separate common marmoset (*Callithrix jacchus*) breeding colonies were defined using 16S rRNA sequencing of rectal swab samples. Study animals originated from two German colonies and a United States colony (JHU). The two German cohorts, previously fed the same diet, were imported into the JHU facility; they were then isolated, transitioned onto JHU diet, and then moved into rooms housing JHU animals. To dissect the contributions of diet and integration in shaping the rectal bacterial community, samples were collected from German origin marmosets upon JHU arrival (baseline), following diet transition (100 d), and following cohousing (390 d). Baseline and 390 d samples were collected from stably maintained JHU marmosets. Bacterial community composition was distinct between all three cohorts at baseline, suggesting that factors other than primary diet confer significant differences between captive populations. Beta-diversity of the animals from the two German colonies converged by 100 d but remained distinct from JHU sample beta-diversity throughout the 390-d study, indicating that diet had greater influence on bacterial community composition than did housing animals within the same room. Our results demonstrate substantial differences in gut bacteria between different captive marmoset colonies, with persistence of these differences following husbandry standardization and housing integration. Goals of rigor and reproducibility in research underscore the need to consider microbial differences between marmosets of diverse origin.

**Importance:** Characterizing gut microbial populations is expected to promote health and enhance research reproducibility in animal studies. As use of common marmosets as animal models of human diseases expands, evaluating the marmoset gut bacterial community will be critical for interpreting research findings, especially as marmosets are prone to gastrointestinal inflammation. In this study, using 16S rRNA sequencing of rectal swab samples, we compared bacterial community among three captive colonies of marmosets at baseline and following importation of cohorts from two of the colonies into the third colony. Diet history had sustained influence on bacterial community composition, while housing the animals within the same room over a period of eight months did not appear to be a major factor. These persistent differences in marmoset gut bacterial community highlight the need for careful consideration of animal origin as a variable in marmoset research studies.

## Introduction

Interactions between commensal microbes and their human and animal hosts have wide-reaching effects on physiology. In the context of animal research, these effects must be considered in the development and maintenance of robust model systems. In particular, the microbiome of the gut comprises the largest and most diverse microbial population in mammals, and characterization of this system will be helpful in understanding and managing natural and experimental disease states in research animals. The common marmoset (*Callithrix jacchus*) is an increasingly utilized translational model within the fields of behavior, neuroscience, nutrition, and aging. However, it is unknown whether and to what extent the gut microbiota differs between marmosets of differing captive origin. Importantly, limited availability of marmosets for biomedical research often requires animals to be obtained from a variety of colonies for inclusion in a single study cohort. The need to maintain genetically diverse breeding pools also prompts transfer of marmosets between colonies, with unknown implications for ongoing research models.^1^

Broadly, environmental factors are a key source of microbial variability among individuals. In one study of the gut bacterial microbiota in multiple marmoset species and hybrid types, the degree of captivity – captive-born vs. wild-born captive vs. wild – had a stronger overall effect on the microbial composition than did host genetics.^2^ Importantly, there is reason to believe that even conditions of captivity that are relatively similar between colonies can yield significantly differential gut microbial community composition. In inbred mice populations originating from different vendors, or arriving at different facilities with standardized husbandry practices, significant gut microbial variation exists and has been shown to impact research outcomes and disease phenotype.^3–7^ Captive care and husbandry for marmosets has been an ongoing challenge in maintaining these arboreal, exudativorous new world primates in captive settings, and practices to date are unstandardized in comparison to those for mice used in research.^8^ Given variation reported in mouse models in stable and changing conditions of captivity, it is probable that (1) the gut microbiota of captive common marmosets varies widely between colonies, and (2) there is a significant degree of gut microbial community plasticity following minor (i.e. captive to captive) environmental change in the common marmoset.

Recent attention to the variation in husbandry practices has underscored the relevance of such variation in research reproducibility.^8,9^ Strikingly, one research group found a 25% reduction in clinically evident recombinant protein-induced experimental autoimmune encephalomyelitis (EAE) in common marmosets fed a yoghurt-based supplemental diet that differed from the standard colony enrichment; the primary diet for both groups was the same. Divergence in the fecal bacterial microbiota (bacteriota) of the two groups was minimal at seven weeks following implementation of varied dietary supplementation, but lower abundance of *Bifidobacterium spp.* and higher abundance of *Collinsella tanakaei* was noted in the traditional supplementation (non-yoghurt-based) group three weeks following EAE induction, suggesting interplay between diet, the microbiota, and immune function in this model.^9^ This is not an isolated example. In humans and other species, the gut microbiome has been widely implicated in development and modulation of the nervous system via the gut-brain axis,^10–16^{O’Mahony, 2015 #1505;Zheng, 2019 #1483;Zhu, 2019 #1474;Hsiao, 2013 #1726} and in progression of both neurodevelopmental and neurodegenerative disease phenotypes.^9,17–19^ Particularly with the emergence of transgenic, induced, and spontaneous common marmoset models of human neurodegenerative diseases including polyglutamine diseases (e.g. Huntington’s disease and spinocerebellar ataxia), Alzheimer’s disease, Parkinson’s disease, schizophrenia, and amyotrophic lateral sclerosis,^20,21^ it is imperative that the research community consider gut-microbiome-brain interactions in the context of these translational models.

In the current study, we used the rectal bacterial microbiota as a representative component of the gut microbiota. Rectal microbial populations correlate highly with other segments of the gut in nonhuman primates.^22^ We examined whether and to what extent (1) the gut bacterial microbiota of marmosets originating from separate captive colonies displayed significant deviation; (2) institutional transfer conferred significant differences in the gut fecal microbiota of these animals; and (3) the gut bacterial microbiota in adult marmosets within a representative, stable captive research colony maintained stability over time. For 390 d following importation of healthy cohorts from an established academic research German colony (HHG) and an established primate research center Germany colony (DPG), we assessed and compared rectal bacterial microbiota of these marmosets and of healthy colony animals from an established United States academic research colony (JHU) of over 120 individuals.

Based on previous work indicating environmental and genetic effects on the gut microbiota in mammalian species,^23,24^ we hypothesized that the gut bacterial communities of the three captive common marmoset populations, as assessed by 16S rRNA sequencing and bioinformatics analysis, would differ. We hypothesized that greater discrepancy would exist between the German-origin (DPG, HHG) and JHU groups than between DPG and HHG at baseline, given (1) the same primary diet fed to DPG and HHG, with a different primary diet fed to JHU, and (2) more recent microbial and genetic transmission between DPG and HHG populations. In evaluating longitudinal changes within each of these three populations, we hypothesized greater change in bacterial community structure of the German-origin groups over time as compared to the JHU group, which underwent no husbandry or diet change. To determine the relative contributions of variables relevant to captive research colonies, correlational and diversity analyses were conducted considering the following host characteristics: sex, age, weight, and housing status (single versus pair- or group-housed).

## Results

### Sequencing Summary

In total, over 6.5 million 16S rRNA gene sequences were obtained across the entire dataset post-quality filtration and denoising. A range of 9,600 – 129,000 filtered sequences per sample were obtained, with a minimum requirement of each sample yielding at least 5000 sequences to be included in downstream diversity analyses. Over 1300 unique amplicon sequence variants (ASVs) were identified across all processed samples.

### Evaluation of Rectal Bacterial Community Composition at Baseline

#### Sample Diversity (alpha diversity) by Facility Origin

There was no difference in gut bacterial species richness between the three cohorts at baseline (observed ASVs, P > 0.05; Figure 2A). There was increased bacterial evenness in the JHU cohort compared to the HHG cohort (Heip’s evenness, p = 0.006; Figure 2B). Bacterial evenness in the DPG cohort was intermediate and did not differ from that of HHG or JHU animals (Heip’s evenness, p > 0.05; Figure 2B).

**Figure 1.**
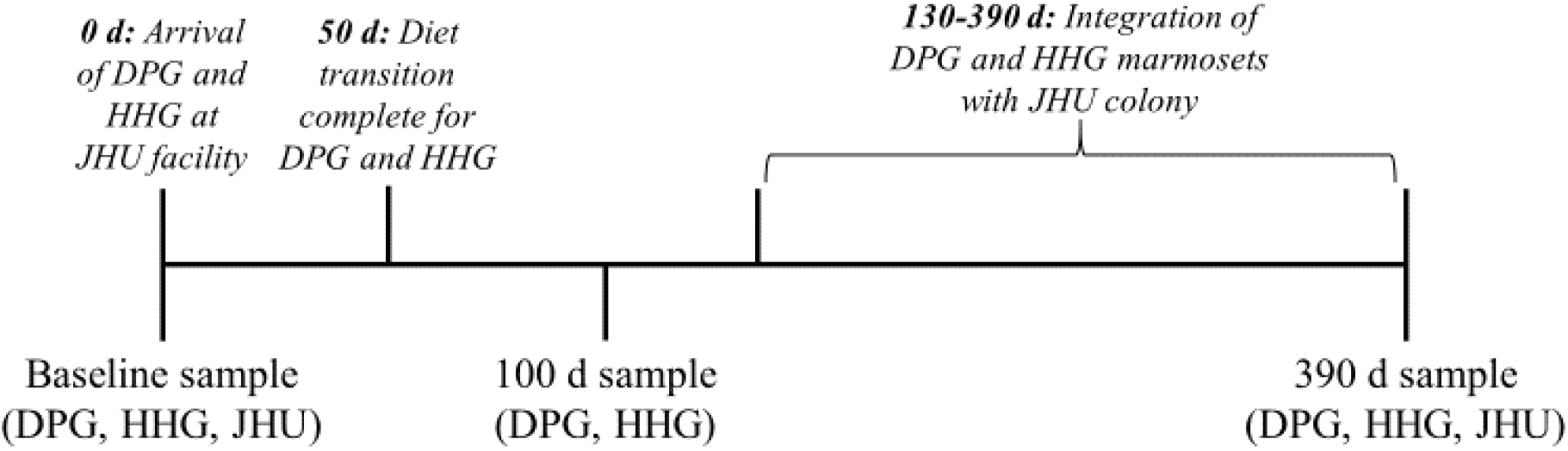
Experimental timeline. Two cohorts of common marmosets from captive research colonies in Germany (DPG, HHG) were imported to the United States and housed at an academic research institution with an existing captive colony (JHU). German-origin marmosets were transitioned from original diet (Mar V3843) onto JHU diet (Callitrichid Diet 5LK6) between days 1 and 50, and were released from quarantine and housed in three rooms containing the entirety of the JHU colony beginning on day 130. Rectal swab samples were collected as indicated at baseline, 100 d, and 390 d.

**Figure 2.**
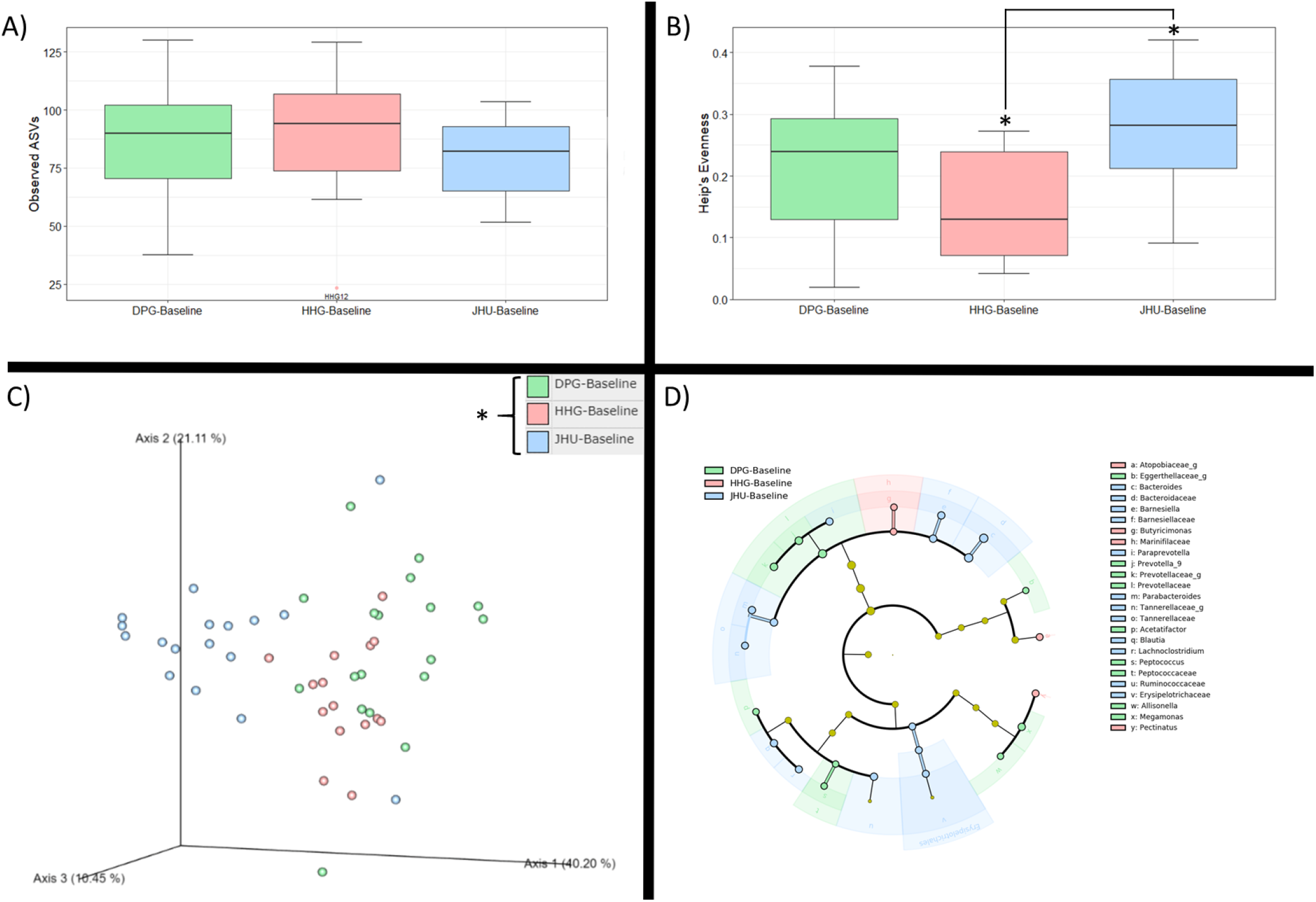
Effect of originating institution on the rectal bacterial community of captive common marmosets. A) Box plot of observed ASVs (alpha-diversity) in the DPG, HHG, and JHU populations at baseline demonstrates no difference between any cohorts. B) Box plot of Heip’s Evenness (alpha-diversity) comparing DPG, HHG, and JHU populations at baseline demonstrates a significant difference between HHG and JHU (p = 0.006). C) PCoA plot with PERMANOVA demonstrates distinct rectal bacterial community structure (beta-diversity) between DPG, HHG, and JHU populations at baseline (p = 0.001). D) Cladogram depicts bacterial taxa that are more abundantly present in DPG (n = 9), HHG (n = 4), and JHU (n = 12) cohorts at baseline. (LDA > 2.5). * = P < 0.05. ASV = amplicon sequence variants. PCoA = principle coordinates analysis. PERMANOVA = permutational analysis of variance. LDA = linear discriminant analysis.

#### Sample Composition (beta diversity) by Facility Origin

Facility origin was associated with significant differences in rectal bacterial community composition (Analysis of similarities: ANOSIM, p = 0.002_DPGvs.HHG,_ p = 0.001_DPGvs.JHU,_ p = 0.001_HHGvs.JHU_; Figure 2C), as well as with observed differences in relative abundance of individual taxa (Figure 2D; Table 1). The majority of DPG samples were dominated by Bacteroidetes (66.7%) or Campylobacterota (27.8%), most HHG samples were dominated by phyla Proteobacteria (53.3%) or Bacteroidetes (40%), and most JHU samples were dominated by Bacteroidetes (77.8%) or Fusobacteria (11.1%) (Table 1). Campylobacterota (or Epsilonbacteraeota) is a recently-proposed phylum comprising members of the class *Epsilonproteobacteria* within the phylum Proteobacteria; genera include *Campylobacter* and *Helicobacter*.^25,26^

**Table 1.** At the baseline and 390 d time points, percentage of common marmosets in the DPG, HHG, and JHU cohorts with rectal bacterial communities dominated by noted bacterial phyla. Campylobacterota is a proposed phylum comprising members of the class *Epsilonproteobacteria* within the phylum Proteobacteria; genera include *Campylobacter* and *Helicobacter*.

| Bacterial Phylum | DPG (%) |  | HHG (%) |  | JHU (%) |  |
| --- | --- | --- | --- | --- | --- | --- |
|  | Baseline<br>(n = 18) | 390 d<br>(n = 16) | Baseline<br>(n = 15) | 390 d<br>(n = 14) | Baseline<br>(n = 15) | 390 d<br>(n = 13) |
| Actinobacteria | 0 | 0 | 0 | 7.1 | 5.6 | 7.1 |
| Bacteroidetes | 66.7 | 56.3 | 40 | 50 | 77.8 | 50 |
| Firmicutes | 0 | 6.3 | 6.7 | 21.4 | 0 | 21.4 |
| Fusobacteria | 0 | 0 | 0 | 21.4 | 11.1 | 21.4 |
| Proteobacteria (traditional classification) | 33.4 | 37.5 | 53.3 | 0 | 5.6 | 0 |
| Proteobacteria (excludes Epsilonproteobacteria / Campylobacterota) | 5.6 | 0 | 53.3 | 0 | 5.6 | 0 |
| Epsilonproteobacteria / Campylobacterota | 27.8 | 37.5 | 0 | 0 | 0 | 0 |

### Evaluation of Rectal Bacterial Community Composition Following Diet Change for DPG and HHG (100 d), and Integration of All Populations (390 d)

#### Sample Diversity over Time

In the DPG cohort, there was higher bacterial richness noted at 390 d compared to 100d (p = 0.01); however, evenness in the DPG cohort did not significantly differ between any time points (Figure 3A-B). There was no change in richness or evenness between baseline and the 390 d time point in HHG marmosets (observed ASVs, p > 0.05; Heip’s evenness, p > 0.05; Figure 3A-B). However, in this cohort there was an increase in bacterial richness between the 100 d and 390 d time points (Observed ASVs, p = 0.003) and an increase in evenness between the baseline and 100 d time points (Heip’s evenness, p = 0.045). In general, trends suggest a nadir in richness and a peak in evenness at the 100 d time point for the HHG cohort.

**Figure 3.**
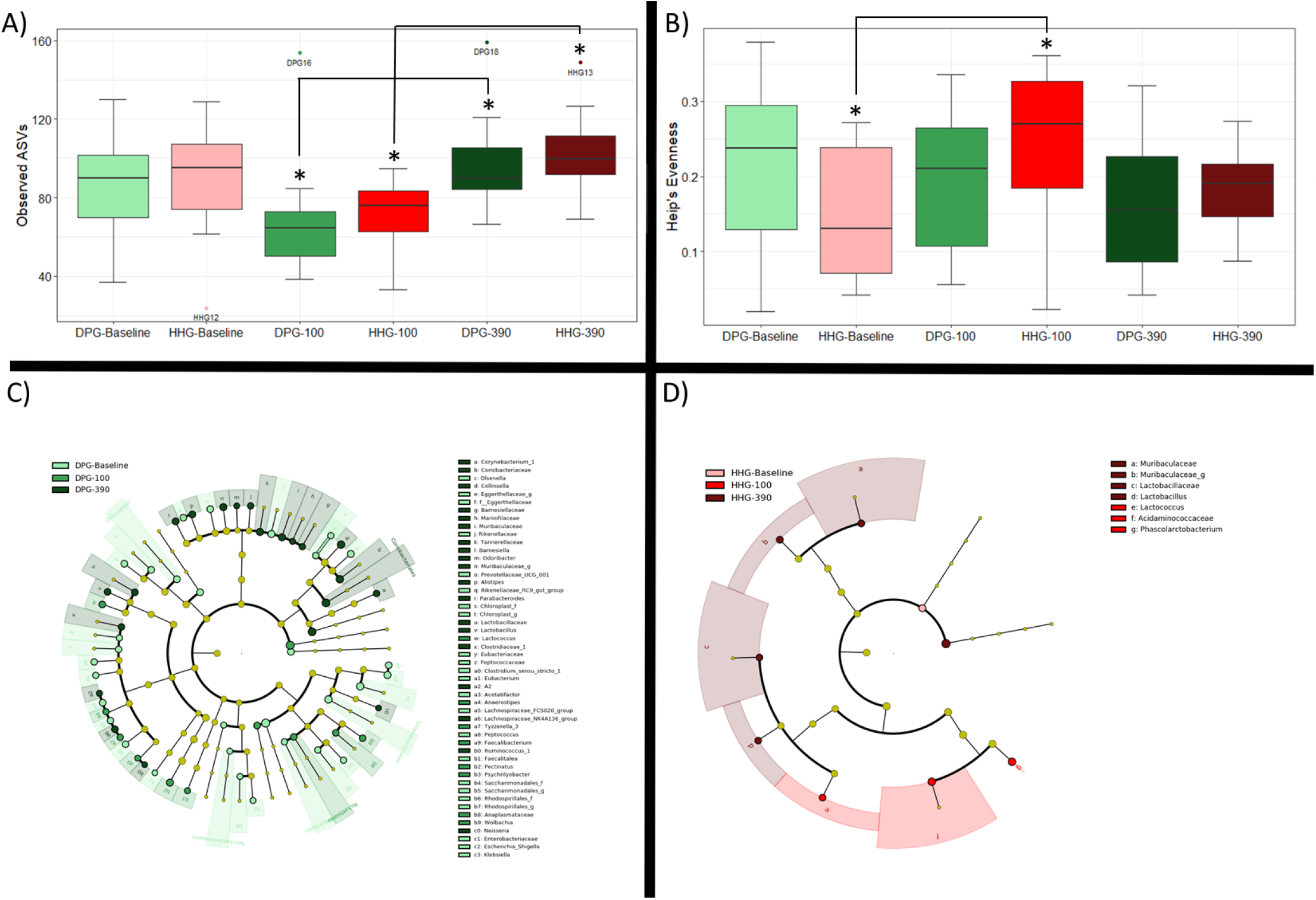
Changes in the rectal bacterial community in imported captive common marmosets (DPG and HHG) from baseline to 390 d. A) Box plot of observed ASVs (alpha-diversity) in the DPG (green) and HHG (red) populations at baseline (lightest color saturation), 100 d (medium), and 390 d (dark). In the HHG and DPG cohorts, observed ASV richness is significantly increased between 100 days post-arrival and 390 d post-arrival (HHG p = 0.003; DPG p = 0.01). B) Box plot of Heip’s evenness (alpha-diversity) in the bacterial community of DPG and HHG populations at baseline, 100 d, and 390 d. In the HHG cohort, there is increased Heip’s evenness from baseline to the 100 day time point (p = 0.045). C) Cladogram depicts bacterial taxa that are more abundantly present at 100 d (n= 3) and 390 d (n = 4) in the DPG cohort (LDA > 2.5). D) Cladogram depicts bacterial taxa that are more abundantly present at baseline (n = 23), 100 d (n = 8), and 390 d (n = 19) in the HHG cohort (LDA > 2.5). * = P < 0.05. ASV = amplicon sequence variants. LDA = linear discriminant analysis.

Following integration with the arrived German cohorts (390 d), the bacterial population of JHU animals decreased in evenness (Heip’s evenness, p = 0.002; Figure 4B). There was a non-significant trend suggesting an increase in richness (Observed ASVs, p > 0.05; Figure 4A).

**Figure 4.**
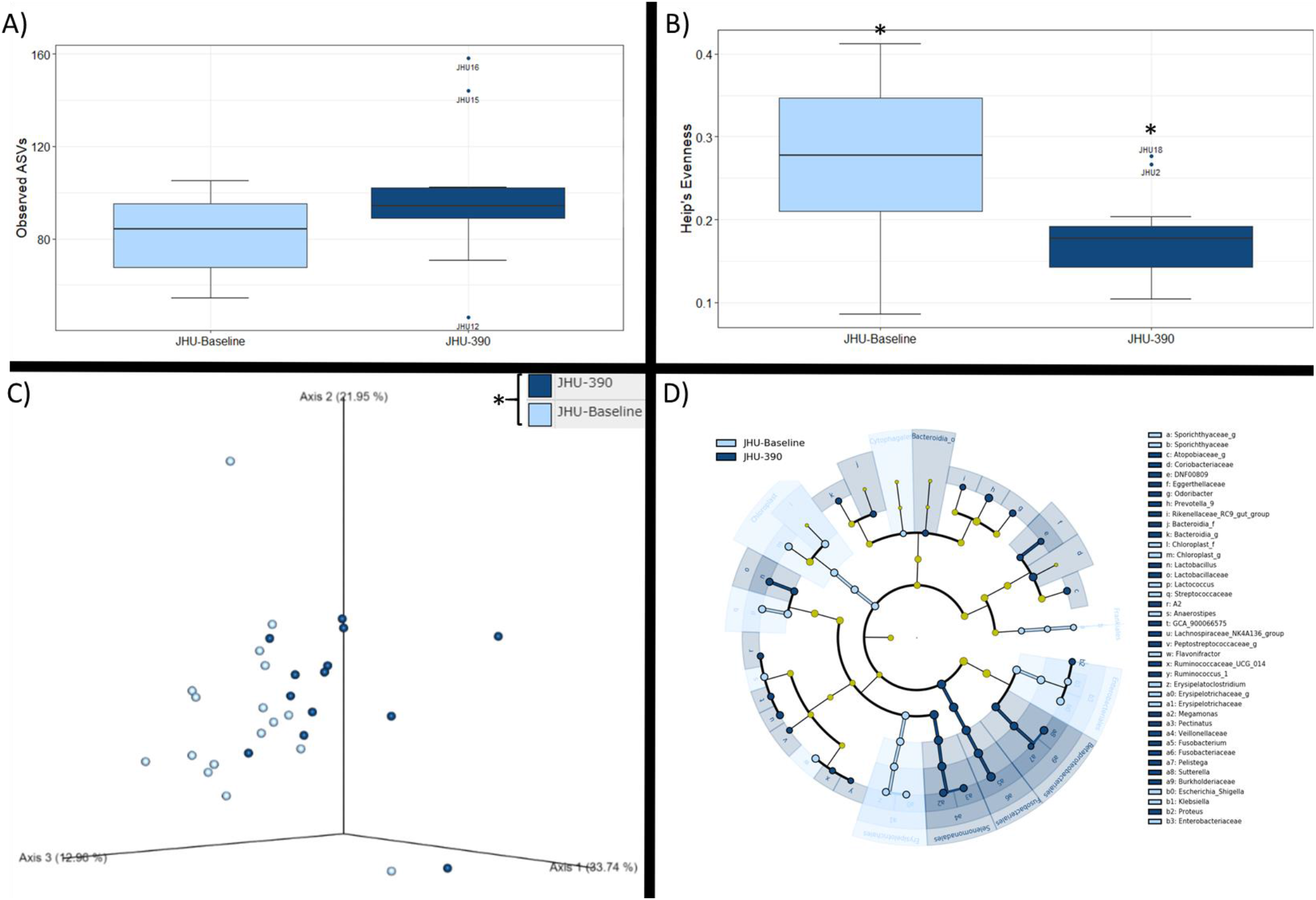
Changes in the rectal bacterial community in a stable population of captive common marmosets (JHU) from baseline to 390 d. A) Box plot of observed ASVs (alpha-diversity) in the JHU cohort shows no change from baseline (light blue) to 390 d (dark blue) (p = 0.09). B) In the JHU cohort, bacterial evenness (Heip’s evenness) is significantly decreased at the 390 d time point compared to baseline (p = 0.002). C) PCoA plot with PERMANOVA demonstrates distinct bacterial community structure (beta-diversity) between baseline and 390 d in the JHU population (p = 0.022). D) Cladogram depicts bacterial taxa that are more abundantly present at baseline (n = 14) and 390 d (n = 26) in the JHU cohort (LDA > 2.5). * = p < 0.05. ASV = amplicon sequence variants. PCoA = principle coordinates analysis. PERMANOVA = permutational analysis of variance. LDA = linear discriminant analysis.

At 390 d, there was no difference between the DPG, HHG, and JHU cohorts in observed species or Heip’s evenness (p > 0.05; Figure 5A-B).

**Figure 5.**
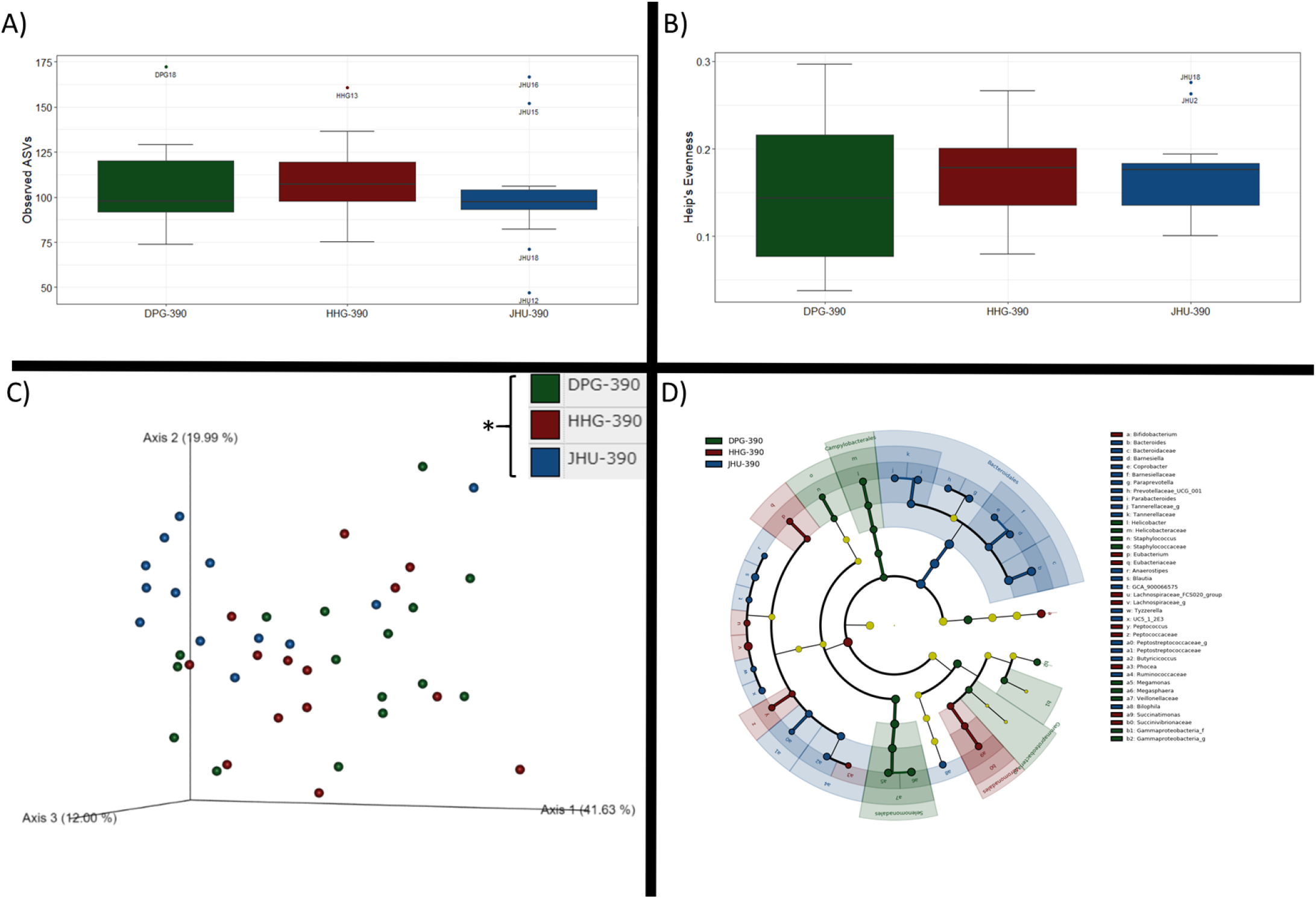
Effect of originating institution (DPG, HHG, JHU) on the rectal bacterial community of common marmosets at 390 d. A,B) Box plots of observed ASVS and Heip’s Evenness (alpha-diversity) in the HHG, DPG, and JHU common marmoset populations at 390 d demonstrate no difference between any cohorts. C) PCoA plot with PERMANOVA demonstrates distinct bacterial community structure (beta-diversity) between HHG, DPG, and JHU populations at 390 d (p = 0.002). D) Cladogram depicts bacterial taxa that are more abundantly present in DPG (n = 9), HHG (n = 10), and JHU (n = 20) cohorts at 390 d. * = p < 0.05. ASV = amplicon sequence variants. PCoA = principle coordinates analysis. PERMANOVA = permutational analysis of variance. LDA = linear discriminant analysis.

#### Sample Composition over Time

Within each of the three cohorts, bacterial community composition differed from baseline to 390 d (ANOSIM, p = 0.011 _JHU_, p = 0.002_HHG_, p = 0.02_DPG_; Figures 3C-D, 4C-D; Table S2). In pairwise comparisons of each of the three time points for DPG and HHG cohorts, significant differences for each population group were observed between every time point, except for between 100 d and 390 d in the DPG cohort (ANOSIM; Table S2).

At 100 d and 390 d, there was no significant difference in bacterial community composition between DPG and HHG cohorts (ANOSIM, p = 0.082_100 d_, p = 0.573_390 d_; Table S3). At 390 d, the bacterial microbiota of DPG and HHG marmosets remained distinct from that of JHU marmosets (ANOSIM; p = 0.006_DPGvs.JHU_, p = 0.004_HHGvs.JHU_; Table S3).

At 390 d, there were persistent differences at the phylum, class, order, and family levels; there were 39 differential taxa, compared to 25 differential taxa at baseline (Linear discriminant analysis: LDA > 2.5; Figures 2D, 5D). At 390 d, most samples from all cohorts were predominated by the phylum Bacteroidetes (DPG: 563%, HHG: 50%, JHU: 76.9%); phyla of secondary predominance included Epsilonbacteraeota (37.5%) for DPG marmosets, Firmicutes (21.4%) and Fusobacteria (21.4%) for HHG marmosets, and Fusobacteria (23.1%) for JHU marmosets (Table 1).

### Evaluation of Study Subject Characteristics

#### Sample Composition by Sex

At 390 d, with all marmoset cohorts grouped for analysis, there was no effect of sex (male; n = 20 vs. Female; n = 23) on overall bacterial community composition (ANOSIM, q = 0.623, p = 0.623; Figure 6C). However, seven bacterial taxa were significantly enriched in male marmosets (LDA > 2.5, Figure 6D).

**Figure 6.**
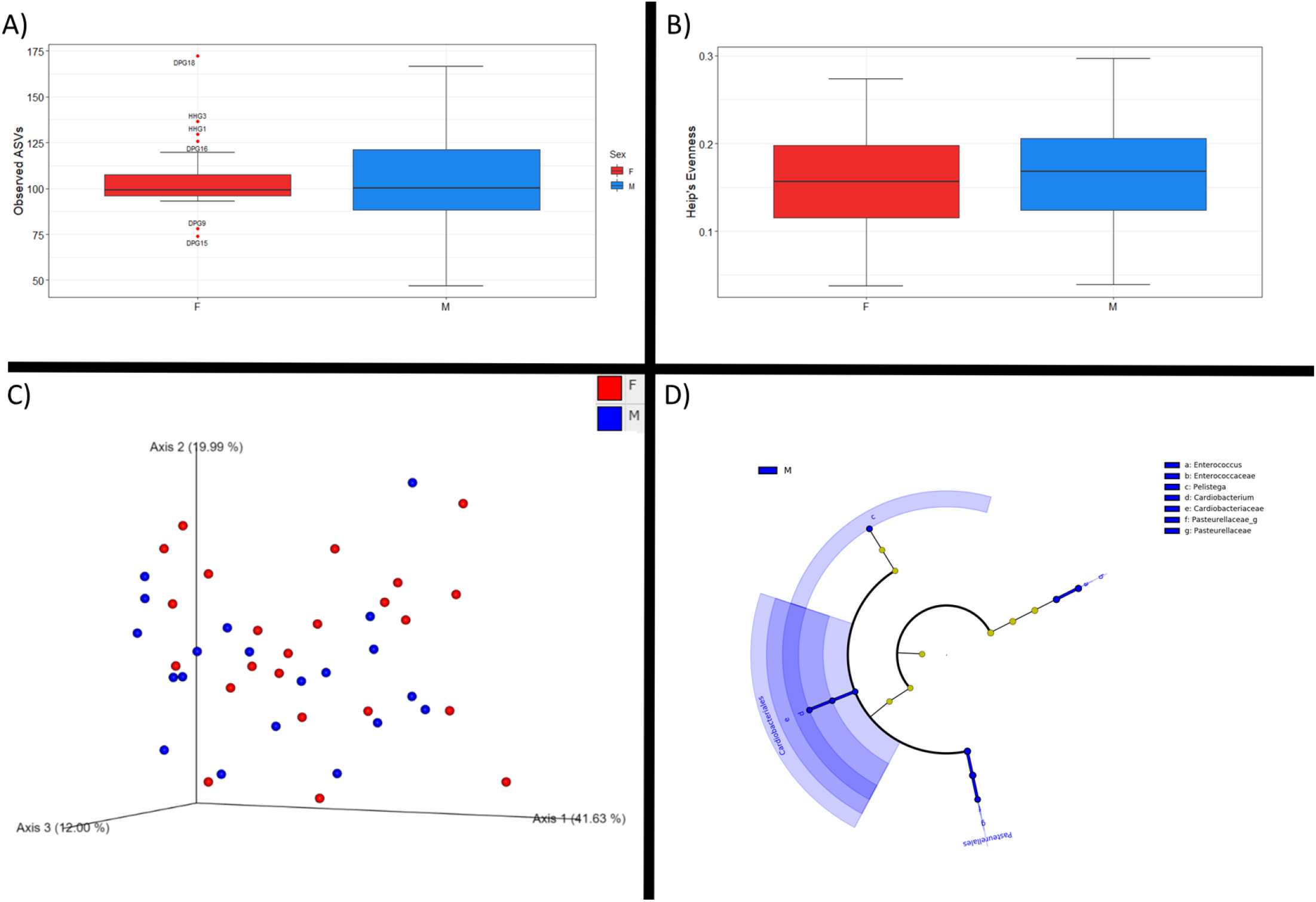
Effect of sex (male vs. female) on the rectal bacterial community of common marmosets at 390 d. Data from DPG, HHG, and JHU cohorts are grouped for analysis. A,B) Box plots of observed ASVS and Heip’s Evenness (alpha-diversity) in male and female marmosets demonstrate no difference between sexes. C) PCoA plot with PERMANOVA demonstrates no difference in bacterial community composition between male and female marmosets (p = 0.623). D) Cladogram depicts bacterial taxa that are more abundantly present in male marmosets (n = 7) (LDA > 2.5). ASV = amplicon sequence variants. PCoA = principle coordinates analysis. PERMANOVA = permutational analysis of variance. LDA = linear discriminant analysis.

#### Sample Composition by Housing Status

At 390 d, with all marmoset cohorts grouped for analysis, there was no effect of housing status (singly vs. pair- or group-housed) on bacterial community composition (ANOSIM, p = 0.38).

#### Sample Composition by Age

At 390 d, with all marmoset cohorts grouped for analysis, two bacterial taxa varied significantly with age; both increased in abundance as animal age increased (Spearman correlation: genus *Phascolarctobacterium*, *r* = 0.380, p = 0.017; family Peptostreptococcaceae *r* = 0.372, p = 0.019; Table S4).

#### Sample Composition by Body Mass

At 390 d, with all marmoset cohorts grouped for analysis, 13 bacterial taxa varied significantly with body mass. Six taxa (*Clostridium sensu stricto*, *Fusobacterium mortiferum*, *Butyricicoccus*, *Cellulosilyticum*, *Enterococcus*, and *Phoceamassiliensis*) correlated positively (Spearman: *r* = 0.322-0.410, p < 0.05; Table S5); 7 taxa (Bifidobacteriaceae, *Bifidobacterium aesculapii*, *Lachnoclostridium*, Muribaculaceae, Atopobiaceae, Peptostreptococcaceae, and *Phascolarctobacterium*) correlated negatively (Spearman: *r* = −0.328- −0.404, p < 0.05; Table S5). There were no significant correlations above the order taxonomic level.

## Discussion

The results of this study emphasize the importance of animal source as a potential variable for research. We found significant baseline differences in the rectal bacterial community composition of two different captive populations (DPG, HHG) originating from the same country, fed the same primary diet, and with limited cross-institutional animal transfer in recent years. Although there were no differences in overall bacterial community structure (beta-diversity) by 100 d following colony integration, significant differences in the presence of certain bacterial taxa (e.g. *Helicobacter jaachi*) persisted for the length of the study. Rectal bacterial differences were even more persistent between populations that had previously been fed different primary diets and had different captive country origin (DPG, HHG vs JHU); bacterial community composition between such populations remained significantly different for over one year. Greater likelihood of genetic similarity between DPG and HHG populations is unlikely to explain these results; previous studies in nonhuman primates have found that environment – diet in particular – appears to be much more important than host genetics in predicting the composition of the gut microbiota.^2,24^

390 d after importation, bacterial community composition remained distinct between the imported population groups (DPG, HHG) and the stable population group (JHU). These results suggest a level of long-term stability of the gut microbiota in captive marmosets when moved between institutions and subjected to diet change. Over the course of the study – while all animals were housed at JHU – the rectal bacterial microbiota of the two German cohorts converged, predominantly between the baseline and 100 d time points. This suggests that diet change and standardization of other environmental parameters (e.g. water source, non-edible enrichment items, air quality) were instrumental in altering the microbiota. Interestingly, though sample composition was not significantly different between the DPG and HHG groups by 100 d, each population retained distinctive microbial signatures throughout the study. DPG marmoset samples, for example, remained uniquely enriched in *Epsilonproteobacteria* (proposed phylum Campylobacterota),^25,26^ specifically *Helicobacter jacchi*,^27^ through 390 d.

Although differences between the German cohorts and the JHU cohort persisted through 390 d, we did not find evidence to attribute this fully to long-term stability in the JHU gut bacterial microbiota. Rather, significant alterations in bacterial community structure developed in the JHU group over time. The magnitude of alteration was surprising, given that animals in the JHU cohort did not undergo any substantial changes in husbandry or care practices other than introduction of two relatively small cohorts, comprising 57 marmosets total, into three adjacent housing rooms containing an existing colony of approximately 120 marmosets. One likely explanation for this alteration is that the introduction of new marmosets induced changes in microbial community structure, either through introduction of novel bacteria or other community-altering flora (e.g. phages, fungi) or via other physiological effects (e.g. stress). Other possibilities include random change over time (i.e. drift), or aging of animals within the cohort.

We conducted correlational analysis in order to evaluate the possibility of age as a confounder in this longitudinal study. In humans, aging has been associated with changes in the gut microbiome; with increased age, there has been demonstrated a lack of diversity in the core bacterial taxa and a loss of microbiota stability.^28–30^ Among adult marmosets, we found minimal impact of age on composition of the gut bacterial population, with two taxa within the phylum Firmicutes, genus *Phascolarctobacterium* and family Peptostreptococcaceae, demonstrating a weak positive correlation with age. These findings differ from those previously reported in marmosets; one recent study of 20 captive common marmosets found increased relative abundance of Proteobacteria and decreased Firmicutes in geriatric marmosets (> 8 years of age) compared to young adults.^31^ Findings in the current study, which demonstrated only two taxa that correlated in abundance with advancing age, may be attributable to the relatively limited diversity of ages represented by our study subjects (2.1 years – 7.82); this range excludes geriatric animals, and represents the large majority of marmosets actively used in research.

In this study population of clinically healthy marmosets, body mass was correlated with abundance of a number of bacterial phyla in the gut microbiota. Within Firmicutes, *Clostridium sensu stricto*, *Butyricicoccus*, *Cellulosilyticum*, *Enterococcus*, and *Phocea massiliensis* were present in increased abundance in common marmosets with higher body mass while *Lachnoclostridium*, Peptostreptococcaceae, and *Phascolarctobacterium* were present in decreased abundance. Within the phylum Bacteroidetes, the family Muribaculaceae was found to weakly negatively correlate with body mass. In humans, an increased Firmicutes:Bacteroidetes ratio has been associated with increased obesity risk;^32^ at the phylum level, we did not identify this in our marmoset population. Abundance of Bifidobacteriaceae (phylum: Actinobacteria), including the species *Bifidobacterium aesculapii*, was negatively correlated with body mass in the present study. Members of this taxonomic family have been associated with anti-obesity effects in mice and humans,^33,34^ and are found in decreased abundance in obese humans.^35,36^ These preliminary findings in healthy marmosets, taken together with supporting evidence in other species, suggests utility in further investigation into the relative abundance and role of Bifidobacteriaceae in the gut of obese marmosets.

We detected no influence of sex on overall bacterial community composition; nonetheless, several individual taxa, including families Cardiobacteriaceae (phylum: Proteobacteria), Pasteurellaceae (phylum: Proteobacteria), and Enterococcaceae (phylum: Firmicutes), were relatively enriched in male marmosets. Studies in mice and humans have demonstrated the presence of sex differences, though such differences appear relatively minor; in studies of inbred mice, for example, strain identity has accounted for greater variation than has sex.^37–39^

The most common and growing use of common marmosets in research is for neuroscience and behavior studies, including development of transgenic animals to model neurodevelopmental and neurodegenerative disease. Scientific literature has broadly established the importance and relevance of the microbiome-brain-gut axis;^12–19^ therefore, particularly given the protracted microbiota differences noted in this study, investigators should be aware of the potentially confounding effects of the gut microbiome in the use of differently originating captive common marmosets in development of research models. Transport of animals from one institution to another, even if marmosets remain on the same diet, may have effects on the microbiome and therefore the research model. Likewise, movement of new animals into an existing colony should be performed with caution and understanding that the gut microbiome of existing colony animals may experience significant alteration. Due to the increasing demand for marmosets in research and limited availability of animals from any single colony, these results will be relevant to most captive research colonies within the United States and elsewhere.

## Materials and Methods

### Study population

Adult animals belonging to three captive populations of common marmosets were examined in the present study. At the time of sample collection, all animals were housed at Johns Hopkins University, an AAALAC-accredited institution; all procedures were approved by the Johns Hopkins University IACUC. Animals were allowed *ad libitum* access to chlorine dioxide-treated water (Quip Labs) delivered in water bottles. Marmosets were fed one of two complete commercial callitrichid diets, detailed below, in addition to daily supplementation with some combination of seeds, dried fruit, wax worms, cheerios, tofu, chickpeas, and fresh fruit (e.g. banana, pineapple, kiwi).

JHU Population: Approximately 120 animals (0 – 12.4 years old) were housed in family groups, in pairs, or singly as part of a long-standing breeding and experimental colony at Johns Hopkins University. Animals actively engaged in experimental use participated in awake behavioral experiments with or without cranial implants, and/or anesthetized neurophysiologic experiments with cranial implants. This population was fed a primary diet of Advanced Protocol® Callitrichid Diet 5LK6 (Purina LabDiet®, St. Louis, MO). This diet contained at least 21% crude protein, at least 7% crude fat, up to 2.5% crude fiber, 12% starch, 35% monosaccharides/disaccharides, 1.1% calcium, 0.64% phosphorus, and 6,600 IU/kg Vitamin D_3_.

HHG population: Seventeen animals (1.7 – 6.8 years old) originating from the University of Düsseldorf (Heinrich-Heine-Universität Düsseldorf, Düsseldorf, Germany) were housed singly or in groups for the duration of import and quarantine. At the original institution, during CDC import quarantine (31 d), and during quarantine at Johns Hopkins University, this population was fed a primary diet of Mar V3843 (ssniff®, Soest, Germany). This diet contained 26% crude protein, 7% fat, 2.5% crude fiber, 1% calcium, 0.7% phosphorus, and 3,000 IU/kg vitamin D_3_.

DPG population: Forty animals (2.8 – 8.7 years old) originating from the German Primate Center (Deutsches Primatenzentrum, Gottingen, Germany) were housed singly for the duration of import and quarantine. At the original institution, during CDC import quarantine (31 d), and during quarantine at Johns Hopkins University, this population was fed a primary diet of Mar V3843 (ssniff®, Soest, Germany).

DPG and HHG populations underwent import and quarantine concurrently; these populations remained quarantined from each other during CDC import quarantine, and were housed in the same room but in original housing groups during the Johns Hopkins University quarantine. The primary diet of populations DPG and HHG was 100% Mar V3843 through experimental day 1 (one day following arrival at JHU), 50% V3843/50% 5LK6 through experimental day 50, and 100% 5LK6 subsequently. The primary diet of the JHU population remained unchanged (100% 5LK6) throughout the study duration.

Adult animals in good clinical health were randomly assigned to this study from the three populations (JHU, DPG, and HHG) following application of inclusion criteria: no history of systemic or chronic disease, 2-8 years of age, ≥ 3 months since antimicrobial use (Table S1). Age and body mass did not differ between groups at baseline (ANOVA; age: F = 0.09, p = 0.91; body mass: F = 0.06, p = 0.94).

### Sampling procedure

JHU animals (n = 18) were sampled once each during a two-month window during Johns Hopkins University quarantine of populations DPG and HHG; no introduction of new animals or changes in husbandry procedures took place during this duration. HHG animals (n = 15) and DPG animals (n = 18) were sampled less than 24 hours following arrival at Johns Hopkins University (baseline). Sampling at the post-diet change time point (100 d) took place for populations 2 and 3; study participants from all three populations were sampled at the 390 d time point, 8 months following full colony integration (Figure 1).

Rectal swab samples were obtained from awake animals using swabs (HydraFlock® Sterile Flocked Collection Devices, Puritan Diagnostics LLC, Guilford, ME) advanced approximately 5 cm from rectum to descending colon; each swab was inserted into a sterile microtube (Fisherbrand® Microcentrifuge Tubes, Fisher Scientific, Hampton, NH), flash-frozen in liquid nitrogen, and stored at −80 °C. All samples were collected between 7 am and 1 pm, prior to feeding.

Out of 135 possible samples, 13 samples (9.6%) were not included in analyses; reasons included failure of subject to meet inclusion criteria at 100 d and/or 390 d (n = 5), death of subject by 390 d (n = 3), transfer of study subject to an outside institution by 390 d (n = 1), unavailability of sample (n = 2), or insufficient DNA yield of sample (n = 2).

### DNA isolation and 16S rRNA gene sequencing analysis

DNA was extracted from thawed samples using the QIAamp PowerFecal DNA Kit (QIAGEN, Hilden, Germany), and quantified by the Qubit® dsDNA HS Assay using a Qubit® 2.0 Fluorometer (Invitrogen, Life Technologies Corporation, Carlsbad, CA). Illumina iTag Polymerase Chain Reactions (PCR) were performed based on the Earth Microbiome Project’s 16S rRNA amplification protocol.^40^ The volume of each reaction was 25 μL and contained (final concentrations) 1X PCR buffer, 0.8 mM dNTP’s, 0.625 U Ex Taq DNA Polymerase (Takara), 0.2 μM 515F barcoded forward primer, 0.2 μM 806R reverse primer and ~10 ng of template DNA per reaction. PCR was carried out on a T100 Thermal Cycler (Bio-Rad, Hercules, CA) using the following cycling conditions: 98 °C for 3 min; then 35 cycles of 98 °C for 1 min, 55 °C for 40 s, and 72 °C for 1 min; final extension was at 72 °C for 10 min; then held at 4 °C. PCR products were visualized on a 2% agarose E-Gel with ethidium bromide (Thermo Fisher Scientific) for bands at ~400 bp. Library pools were size verified using the Fragment Analyzer on the ABI3730 and were quantified with a KAPA Library quantification kit (Kapa Biosystem, Wilmington, MA, USA). After dilution with EBT (Illumina) to a final concentration of 2 nM containing 15% PhiX V3 library control (Illumina, San Diego, CA, USA), the library pools were denatured for 5 min in an equal volume of 0.2M NaOH, then further diluted to 8 pM in HT1 buffer (Illumina) and were sequenced using an Illumina MiSeq V2 500 cycle kit cassette with 16S rRNA library sequencing primers set for 250 base pair, paired-end reads. Overall sequencing run performance was evaluated by determining whether the sequencing run met the Illumina specifications for quality scores and data output. Actual run performance varied based on sample type, quality, and clusters passing filter. Specifications are based on the Illumina PhiX control library at supported cluster densities.

#### Bioinformatic Analysis

Paired-end sequences were imported into the DADA2 pipeline for quality filtration, denoising, and chimera removal.^41^ Two hundred fifty base pair reads were filtered at a maximum expected error of 1.0, with forward reads truncated at a length of 218, and reverse at a length of 179. Trimmed sequences were then denoised, merged and underwent chimera removal within DADA2.^42^ Taxonomy was then assigned against the SILVA database.^43^ ASVs were tabulated for use in downstream taxonomic summary and analysis. A reformatted ASV-table was converted to BIOM format for use with QIIME.^44^

Alpha diversity box plots were generated within the *QIIME2* sequence analysis package using an unrarified taxonomy table.^45^ Nonparametric t-test comparisons between experimental groups were made considering the *Observed ASV Richness and Heip’s Evenness.* Cumulative Sum Scaled (CSS) normalized counts of ASVs were imported into QIIME2 after which Principal Coordinates Analysis (PCoA) plots were generated from calculated weighted UniFrac distance matrices. Significance of clustering between samples within all PCoA plots was assessed using the ANOSIM test statistic. For Linear Discriminant Analysis Effect Size (LEfSe) biomarker analysis, relative abundances of bacterial taxa were multiplied by 1 million and formatted as described in Segata et al.^46^ Alpha levels of 0.05 were used for both the Kruskal–Wallis and pairwise Wilcoxon tests. Spearman’s correlational analyses were conducted within QIIME-1.9.1, in which species level taxa summaries were correlated against select continuous metadata variables. ANOVA tests were performed using VassarStats (vassarstats.net) to determine whether key characteristics (i.e. age, body mass) differed between experimental groups at study initiation; there were no differences between groups (p > 0.05). Functional predictions were generated using the PICRUSt2 informatics package from generated ASVs.^47^ Predictions were summarized at the pathway level (L3) and were subject to downstream LEfSe biomarker analysis.

### Data Availability

Sequencing data has been submitted to the NCBI Sequence Read Archive with BioProject ID PRJNA659800.

## Acknowledgments

The project described was supported by the Grants for Laboratory Animal Science (GLAS) from the American Association for Laboratory Animal Science, NIH grant T32 OD011089, and Research Animal Resources at Johns Hopkins University School of Medicine. The funders had no role in study design, data collection or interpretation, or the decision to submit the work for publication. The authors thank Drs. Caroline Garrett and Jessica Izzi for veterinary oversight and consultation throughout the project. We extend special thanks to Dr. Xiaoqin Wang and his research team for generously sharing use of their animals, supported by NIH grants U24MH123423, DC003180, DC005808, and DC014503 (PI: Wang).

## Supplemental Tables

**Supplemental Table 1.**
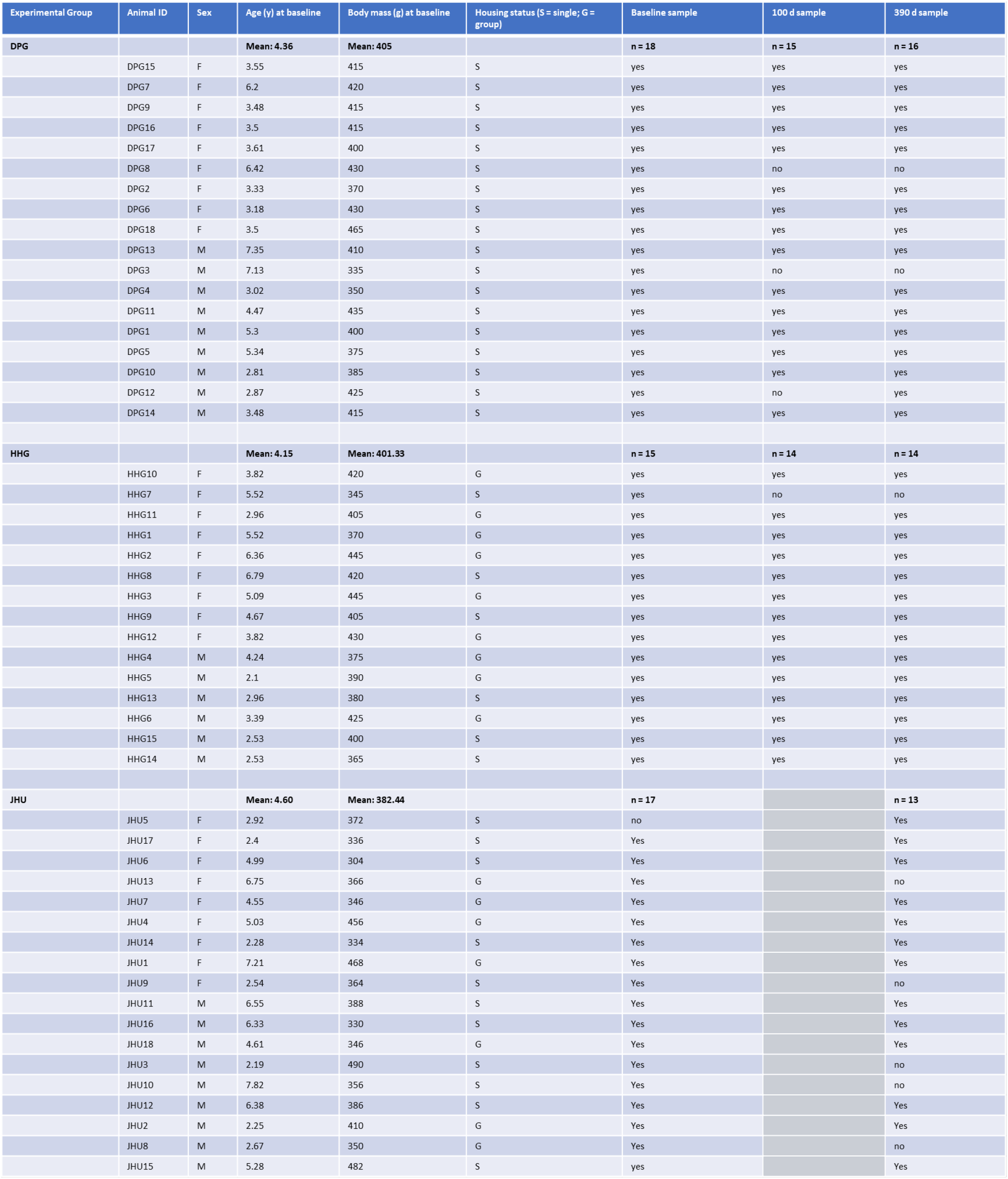
Study groups and subject characteristics. Inclusion criteria: No history of systemic or chronic disease, between 2 and 8 years of age, ≥ 3 months since antimicrobial use.

| Experimental Group | Animal ID | Sex | Age (y) at baseline | Body mass (g) at baseline | Housing status (S = single; G = group) | Baseline sample | 100 d sample | 390 d sample |
| --- | --- | --- | --- | --- | --- | --- | --- | --- |
| DPG |  |  | Mean: 4.36 | Mean: 405 |  | n = 18 | n = 15 | n = 16 |
|  | DPG15 | F | 3.55 | 415 | S | yes | yes | yes |
|  | DPG7 | F | 6.2 | 420 | S | yes | yes | yes |
|  | DPG9 | F | 3.48 | 415 | S | yes | yes | yes |
|  | DPG16 | F | 3.5 | 415 | S | yes | yes | yes |
|  | DPG17 | F | 3.61 | 400 | S | yes | yes | yes |
|  | DPG8 | F | 6.42 | 430 | S | yes | no | no |
|  | DPG2 | F | 3.33 | 370 | S | yes | yes | yes |
|  | DPG6 | F | 3.18 | 430 | S | yes | yes | yes |
|  | DPG18 | F | 3.5 | 465 | S | yes | yes | yes |
|  | DPG13 | M | 7.35 | 410 | S | yes | yes | yes |
|  | DPG3 | M | 7.13 | 335 | S | yes | no | no |
|  | DPG4 | M | 3.02 | 350 | S | yes | yes | yes |
|  | DPG11 | M | 4.47 | 435 | S | yes | yes | yes |
|  | DPG1 | M | 5.3 | 400 | S | yes | yes | yes |
|  | DPG5 | M | 5.34 | 375 | S | yes | yes | yes |
|  | DPG10 | M | 2.81 | 385 | S | yes | yes | yes |
|  | DPG12 | M | 2.87 | 425 | S | yes | no | yes |
|  | DPG14 | M | 3.48 | 415 | S | yes | yes | yes |
| HHG |  |  | Mean: 4.15 | Mean: 401.33 |  | n = 15 | n = 14 | n = 14 |
|  | HHG10 | F | 3.82 | 420 | G | yes | yes | yes |
|  | HHG7 | F | 5.52 | 345 | S | yes | no | no |
|  | HHG11 | F | 2.96 | 405 | G | yes | yes | yes |
|  | HHG1 | F | 5.52 | 370 | G | yes | yes | yes |
|  | HHG2 | F | 6.36 | 445 | G | yes | yes | yes |
|  | HHG8 | F | 6.79 | 420 | S | yes | yes | yes |
|  | HHG3 | F | 5.09 | 445 | G | yes | yes | yes |
|  | HHG9 | F | 4.67 | 405 | S | yes | yes | yes |
|  | HHG12 | F | 3.82 | 430 | G | yes | yes | yes |
|  | HHG4 | M | 4.24 | 375 | G | yes | yes | yes |
|  | HHG5 | M | 2.1 | 390 | G | yes | yes | yes |
|  | HHG13 | M | 2.96 | 380 | S | yes | yes | yes |
|  | HHG6 | M | 3.39 | 425 | G | yes | yes | yes |
|  | HHG15 | M | 2.53 | 400 | S | yes | yes | yes |
|  | HHG14 | M | 2.53 | 365 | S | yes | yes | yes |
| JHU |  |  | Mean: 4.60 | Mean: 382.44 |  | n = 17 |  | n = 13 |
|  | JHU5 | F | 2.92 | 372 | S | no |  | Yes |
|  | JHU17 | F | 2.4 | 336 | S | Yes |  | Yes |
|  | JHU6 | F | 4.99 | 304 | S | Yes |  | Yes |
|  | JHU13 | F | 6.75 | 366 | G | Yes |  | no |
|  | JHU7 | F | 4.55 | 346 | G | Yes |  | Yes |
|  | JHU4 | F | 5.03 | 456 | G | Yes |  | Yes |
|  | JHU14 | F | 2.28 | 334 | S | Yes |  | Yes |
|  | JHU1 | F | 7.21 | 468 | G | Yes |  | Yes |
|  | JHU9 | F | 2.54 | 364 | S | Yes |  | no |
|  | JHU11 | M | 6.55 | 388 | S | Yes |  | Yes |
|  | JHU16 | M | 6.33 | 330 | S | Yes |  | Yes |
|  | JHU18 | M | 4.61 | 346 | G | Yes |  | Yes |
|  | JHU3 | M | 2.19 | 490 | S | Yes |  | no |
|  | JHU10 | M | 7.82 | 356 | S | Yes |  | no |
|  | JHU12 | M | 6.38 | 386 | S | Yes |  | Yes |
|  | JHU2 | M | 2.25 | 410 | G | Yes |  | Yes |
|  | JHU8 | M | 2.67 | 350 | G | Yes |  | no |
|  | JHU15 | M | 5.28 | 482 | S | yes |  | Yes |

**Supplemental Table 2.** Within-cohort comparisons of sample diversity (beta diversity) over time. FDR corrected p values.

| Experimental Group | p value |  |  |
| --- | --- | --- | --- |
|  | Baseline vs. 390 d | Baseline vs. 100 d | 100 d vs. 390 d |
| DPG | 0.002* | 0.007* | 0.057 |
| HHG | 0.020* | 0.002* | 0.021* |
| JHU | 0.011* |  |  |

**Supplemental Table 3.** Between-cohort comparisons of sample diversity (beta diversity) at each experimental time point. FDR corrected p values.

| Experimental Time Point | p value |  |  |
| --- | --- | --- | --- |
|  | DPG vs. HHG | DPG vs. JHU | HHG vs. JHU |
| Baseline | 0.005* | 0.002* | 0.002* |
| 100 d | 0.082 |  |  |
| 390 d | 0.573 | 0.006* | 0.004* |

**Supplemental Table 4.**
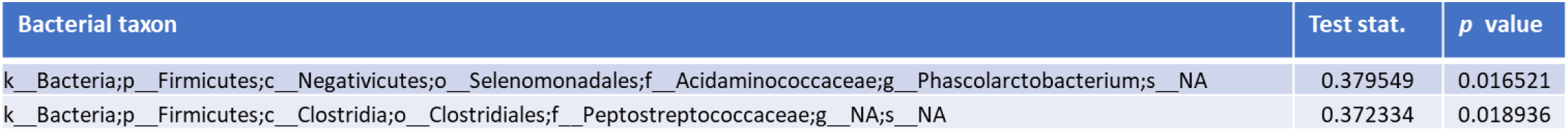
Correlational analysis of rectal bacterial community characteristics associated with marmoset age at 390 d. Spearman correlation detected age-associated alterations in two low-level taxa within the study population (HHG, DPG, and JHU) at 390 d. A positive test statistic indicates positive correlation with increasing age.

| Bacterial taxon | Test stat. | p value |
| --- | --- | --- |
| k_Bacteria;p_Firmicutes;c_Negativicutes;o_Selenomonadales;f_Acidaminococcaceae;g_Phascolarctobacterium;s_NA | 0.379549 | 0.016521 |
| k_Bacteria;p_Firmicutes;c_Clostridia;o_Clostridiales;f_Peptostreptococcaceae;g_NA;s_NA | 0.372334 | 0.018936 |

**Supplemental Table 5.** Correlational analysis of rectal bacterial community characteristics associated with marmoset body mass at 390 d. Spearman correlation detected body mass-associated alterations in 13 low-level taxa within the study population (HHG, DPG, and JHU) at 390 d. A positive test statistic indicates positive correlation with increasing body mass, and a negative test statistic indicates negative correlation.

| Bacterial taxon | Test stat. | p value |
| --- | --- | --- |
| k_Bacteria;p_Firmicutes;c_Clostridia;o_Clostridiales;f_Clostridiaceae_1;g_Clostridium_sensu_stricto_1;s_NA | 0.410456 | 0.00987 |
| k_Bacteria;p_Actinobacteria;c_Actinobacteria;o_Bifidobacteriales;f_Bifidobacteriaceae;g_NA;s_NA | -0.4043 | 0.01119 |
| k_Bacteria;p_Fusobacteria;c_Fusobacteriia;o_Fusobacteriales;f_Fusobacteriaceae;g_Fusobacterium;s_mortiferum | 0.401872 | 0.011751 |
| k_Bacteria;p_Firmicutes;c_Clostridia;o_Clostridiales;f_Ruminococcaceae;g_Butyricoccus;s_NA | 0.401227 | 0.011904 |
| k_Bacteria;p_Actinobacteria;c_Actinobacteria;o_Bifidobacteriales;f_Bifidobacteriaceae;g_Bifidobacterium;s_aesculapii | -0.39686 | 0.012981 |
| k_Bacteria;p_Firmicutes;c_Clostridia;o_Clostridiales;f_Lachnospiraceae;g_Lachnoclostridium;s_NA | -0.38306 | 0.016943 |
| k_Bacteria;p_Firmicutes;c_Clostridia;o_Clostridiales;f_Lachnospiraceae;g_Cellulosilyticum;s_NA | 0.377586 | 0.018768 |
| k_Bacteria;p_Bacteroidetes;c_Bacteroidia;o_Bacteroidales;f_Muribaculaceae;g_NA;s_NA | -0.34853 | 0.03139 |
| k_Bacteria;p_Actinobacteria;c_Coriobacteriia;o_Coriobacteriales;f_Atopobiaceae;g_NA;s_NA | -0.34737 | 0.032009 |
| k_Bacteria;p_Firmicutes;c_Clostridia;o_Clostridiales;f_Peptostreptococcaceae;g_NA;s_NA | -0.34482 | 0.033409 |
| k_Bacteria;p_Firmicutes;c_Bacilli;o_Lactobacillales;f_Enterococcaceae;g_Enterococcus;s_NA | 0.341709 | 0.03518 |
| k_Bacteria;p_Firmicutes;c_Negativicutes;o_Selenomonadales;f_Acidaminococcaceae;g_Phascolarctobacterium;s_NA | -0.32801 | 0.043908 |
| k_Bacteria;p_Firmicutes;c_Clostridia;o_Clostridiales;f_Ruminococcaceae;g_Phocaea;s_massiliensis | 0.322189 | 0.0481 |

